# Cingulo-Opercular Control Network Supports Disused Motor Circuits in Standby Mode

**DOI:** 10.1101/2020.09.03.275479

**Authors:** Dillan J. Newbold, Evan M. Gordon, Timothy O. Laumann, Nicole A. Seider, David F. Montez, Sarah J. Gross, Annie Zheng, Ashley N. Nielsen, Catherine R. Hoyt, Jacqueline M. Hampton, Mario Ortega, Babatunde Adeyemo, Derek B. Miller, Andrew N. Van, Scott Marek, Bradley L. Schlaggar, Alexandre R. Carter, Benjamin P. Kay, Deanna J. Greene, Marcus E. Raichle, Steven E. Petersen, Abraham Z. Snyder, Nico U.F. Dosenbach

## Abstract

Whole-brain resting-state functional MRI (rs-fMRI) during two weeks of limb constraint revealed that disused motor regions became more strongly connected to the cingulo-opercular network (CON), an executive control network that includes regions of the dorsal anterior cingulate cortex (dACC) and insula (1). Disuse-driven increases in functional connectivity (FC) were specific to the CON and somatomotor networks and did not involve any other networks, such as the salience, frontoparietal, or default mode networks. Censoring and modeling analyses showed that FC increases during casting were mediated by large, spontaneous activity pulses that appeared in the disused motor regions and CON control regions. During limb constraint, disused motor circuits appear to enter a standby mode characterized by spontaneous activity pulses and strengthened connectivity to CON executive control regions.

**Significance:** Many studies have examined plasticity in the primary somatosensory and motor cortex during disuse, but little is known about how disuse impacts the brain outside of primary cortical areas. We leveraged the whole-brain coverage of resting-state functional MRI (rs-fMRI) to discover that disuse drives plasticity of distant executive control regions in the cingulo-opercular network (CON). Two complementary analyses, pulse censoring and pulse addition, demonstrated that increased functional connectivity between the CON and disused motor regions was driven by large, spontaneous pulses of activity in the CON and disused motor regions. These results point to a previously unknown role for the CON in supporting motor plasticity and reveal spontaneous activity pulses as a novel mechanism for reorganizing the brain’s functional connections.

## Introduction

Disuse is a powerful paradigm for inducing plasticity that has uncovered key organizing principles of the human brain (2–5). Monocular deprivation revealed that multiple afferent inputs can compete for representational territory in the primary visual cortex (2). Competition between afferents also shapes the somatomotor system, and manipulations such as peripheral nerve deafferentation, whisker trimming, and limb constraint all drive plasticity in the primary somatosensory and motor cortex (3–5). Most plasticity studies to date have used focal techniques, such as microelectrode recordings, to study local changes in brain function. As a result, little is known about how behavior and experience shape the brain-wide functional networks that support complex cognitive operations (6).

The brain is composed of networks of regions that cooperate to perform specific cognitive operations (6–9). These functional networks show synchronized spontaneous activity while the brain is at rest, a phenomenon known as resting state functional connectivity (FC) (10–12). FC can be measured non-invasively in humans using resting-state functional MRI (rs-fMRI). Whole-brain rs-fMRI has been used to parse the brain into canonical functional networks (13, 14), including visual, auditory and somatomotor networks (15, 16); ventral and dorsal attention networks (9, 17); a default mode network with roles in internally directed cognition and episodic memory (8, 12); a salience network thought to assess the homeostatic relevance of external stimuli (18); a frontoparietal control network supporting error-processing and moment-to-moment adjustments in behavior (1, 19, 20); and a cingulo-opercular control network (CON), which maintains executive control during goal-directed behavior (1, 19, 21).

A more recent advance in human neuroscience has been the recognition of individual variability in network organization (22–25). Most early rs-fMRI studies examined central tendencies in network organization using group-averaged FC measurements (11, 13, 14). Recent work has demonstrated that functional networks can be described in an individual-specific manner if sufficient rs-fMRI data are acquired, an approach termed Precision Functional Mapping (PFM) (22, 23, 26–30). PFM respects the unique functional anatomy of each person and avoids averaging together functionally distinct brain regions across individuals.

Somatomotor circuits do not function in isolation. Action selection and motor control are thought to be governed by complex interactions between the somatomotor network and control networks, including the CON (1). Here, we leveraged the whole-brain coverage of rs-fMRI and the statistical power of PFM to examine disuse-driven plasticity throughout the human brain.

## Results

### Disuse Caused Both Increases and Decreases in Functional Connectivity

Three adult participants (“Cast1”, “Cast2” and “Cast3”) were scanned at the same time every day for 42-64 consecutive days (30 minutes of rs-fMRI/day) before, during and after two weeks of dominant upper extremity casting (31). To examine disuse-driven plasticity beyond somatomotor networks, we first generated seed maps showing FC between L-SM1_ue_ and all other cortical regions before, during and after casting (Fig. 1*A*, S1). Prior to casting (Pre), L-SM1_ue_ was strongly connected to the remainder of the somatomotor cortex (pre- and post-central gyri), especially homotopic R-SM1_ue_. During the cast period (Cast), L-SM1_ue_ showed a new pattern of connectivity, with strong connections to bilateral secondary somatosensory cortex (SII), dorsal anterior cingulate cortex (dACC), and pre- and post-central sulci. After casting (Post), L-SM1_ue_ FC returned to baseline. Changes in FC during casting (Cast – Pre) and recovery (Post – Cast) were examined using sets of previously published individual-specific cortical parcels (22, 31). Casting caused decreased L-SM1_ue_ connectivity with the remainder of the somatomotor cortex, as well as increased connectivity with the bilateral SII, dACC, and pre- and post-central sulci (Fig. 1*B*, S1). FC changes after cast removal were strongly negatively correlated with changes during casting (Cast1: spatial correlation, r = −0.56, Cast2: r = −0.95, Cast3: r = −0.32; all participants: P < 0.001), indicating that most effects reversed after cast removal (Fig. 1*B*, S1).

**Fig. 1.**
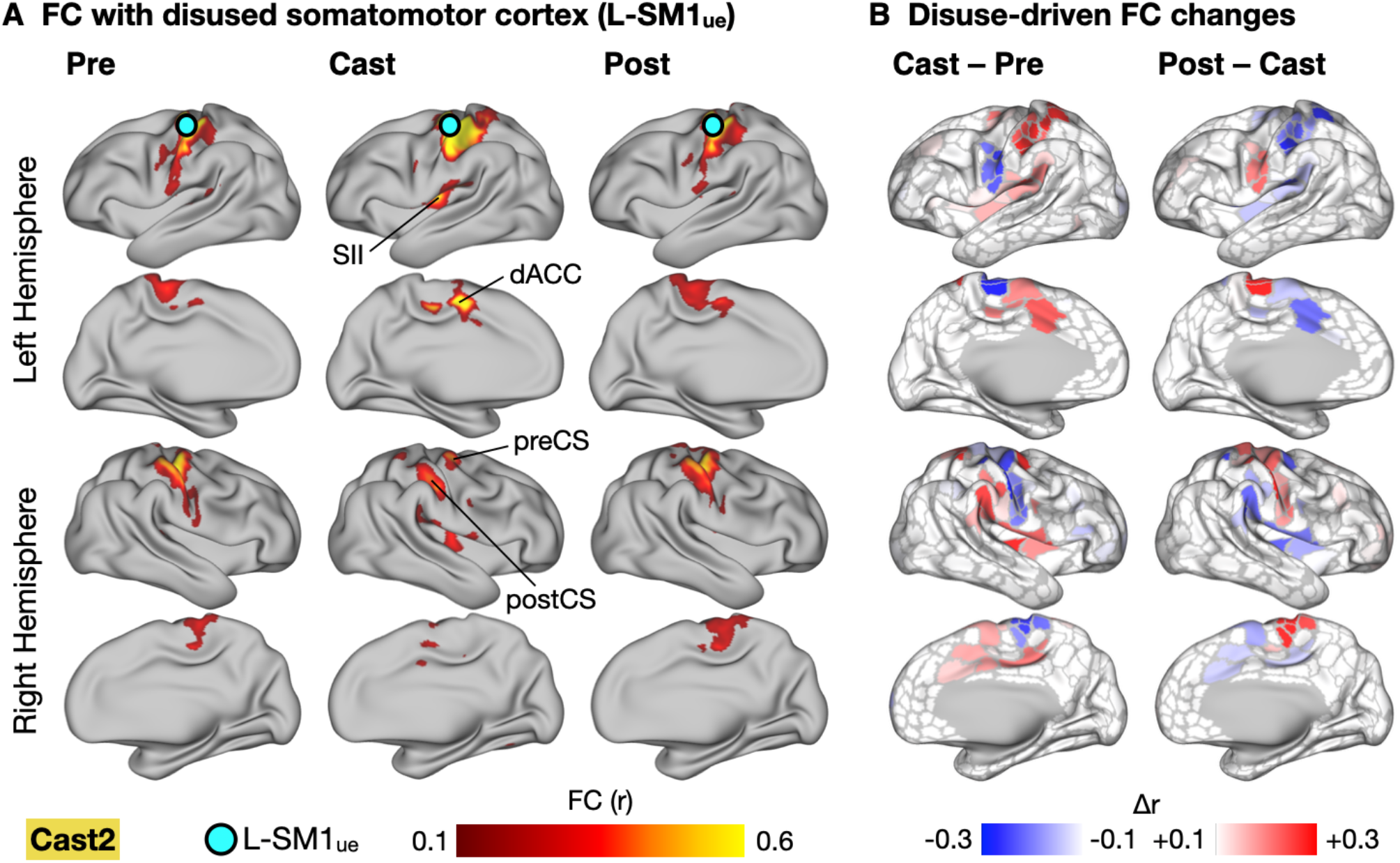
Cast-driven Changes in Functional Connectivity (FC) of Disused Somatomotor Cortex (L-SM1_ue_). (A) Seed maps showing FC between L-SM1_ue_ and the remainder of the cerebral cortex in an example participant (Cast2; remaining participants, Fig. S1). Seed maps were averaged across sessions before, during and after casting (Pre, Cast, Post). Labels indicate secondary somatosensory cortex (SII), dorsal anterior cingulate cortex (dACC), and pre- and post-central sulci (preCS and postCS). (B) Subtraction maps showing changes in FC during casting (Cast – Pre) and recovery (Post – Cast). Differences are shown on a set of previously published individual-specific cortical parcels (22, 31).

### Disuse-driven Functional Connectivity Changes Were Highly Focal in Network Space

The brain’s functional network organization can be visualized using spring-embedded graphs, which treat functional connections as spring forces, positioning more strongly connected nodes closer to one another (Fig. 2*A*, S2). While L-SM1_ue_ FC increases were distributed in anatomical space, including regions on the medial and lateral surfaces of both hemispheres, nearly all of these changes mapped onto a single cluster in network space, the CON (Fig. 2*B*, S2). The network focality of L-SM1_ue_ FC increases was quantified by examining the distribution of the greatest FC increases (top 5%) across the canonical functional networks (Fig. 2*B*). In all participants, the CON showed more FC increases than expected by chance (P < 0.001 for all participants). Decreases in L-SM1_ue_ FC were also localized in network space (Fig. 2*C*, S2). Most of the regions showing decreased FC with L-SM1_ue_ during casting belonged to the somatomotor networks (ue-SMN, le-SMN, face-SMN and premotor). All participants showed more FC decreases than expected by chance in the upper-extremity somatomotor network (ue-SMN; P < 0.001 for all participants). Additionally, two participants (Cast2 and Cast3) showed significant FC decreases in other somatomotor networks, including the lower-extremity and face somatomotor networks (le-SMN and f-SMN) and the premotor network.

**Fig. 2.**
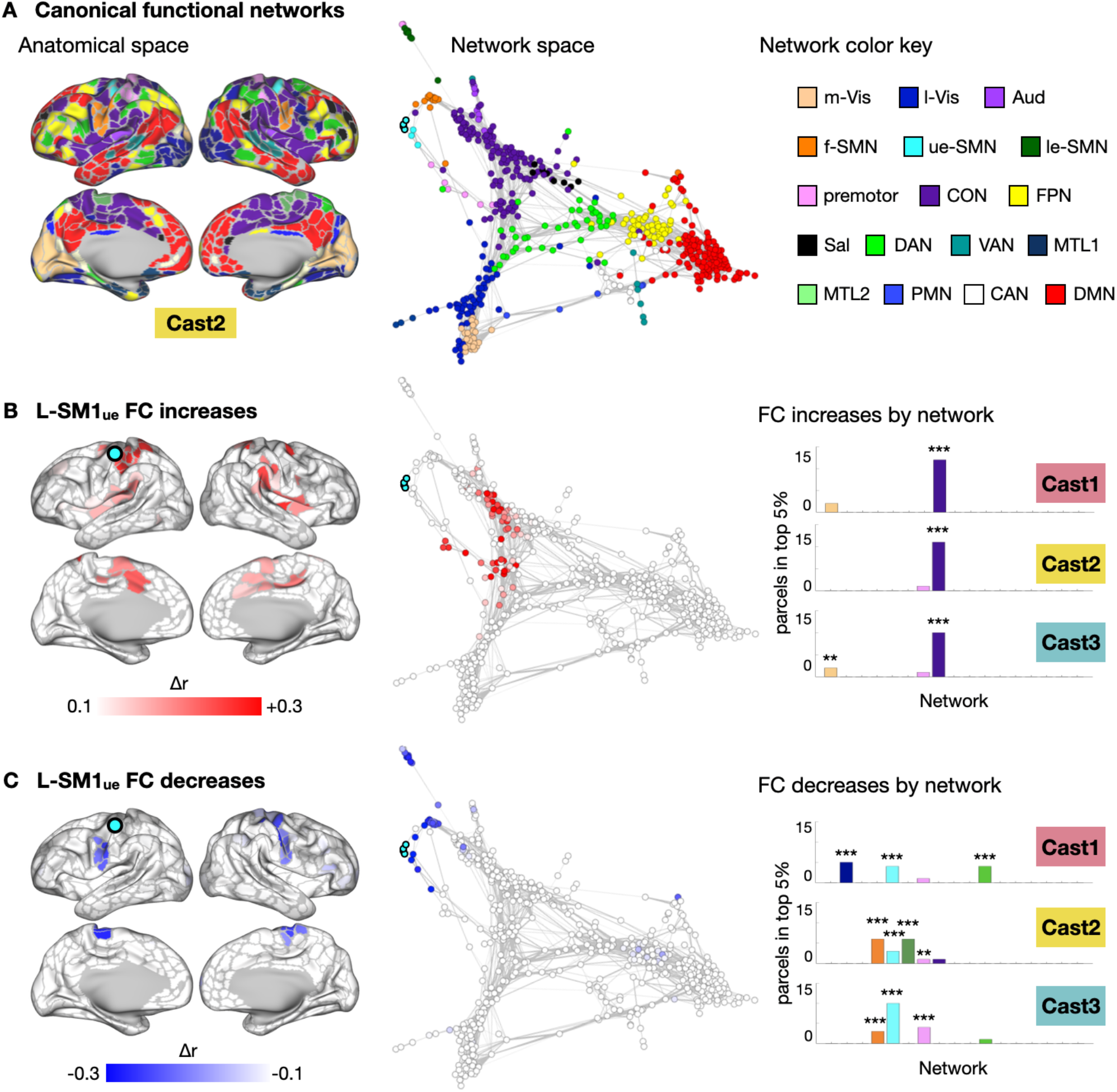
Disused Somatomotor Cortex (L-SM1_ue_) Functional Connectivity (FC) Changes in Network Space. (A) Map of canonical functional networks in an example participant (Cast2; remaining participants, Fig. S1). *Left:* Networks shown in anatomical space (inflated cortical surfaces). *Middle:* Same map as *Top*, shown in network space (spring-embedded graphs based on Pre scans). *Right:* Network color key: m-Vis: medial visual; l-Vis: lateral visual; Aud: auditory; f-SMN: face somatomotor; ue-SMN: upper-extremity somatomotor; le-SMN: lower-extremity somatomotor; CON: cingulo-opercular; FPN: fronto-parietal; Sal: salience; DAN: dorsal attention; VAN: ventral attention; MTL: medial temporal lobe; PMN: parietal memory; CAN: context association; DMN: default mode. (B) *Left:* Subtraction maps showing disuse-driven decreases in functional connectivity (FC) with left somatomotor cortex (L-SM1_ue_) in anatomical space. *Middle:* Same map as *Top*, shown in network space. *Right:* Distribution of top 5% FC decreases across all networks. **P < 0.01; ***P < 0.001; FDR < 0.05. (C) Subtraction maps showing disuse-driven increases in FC with L-SM1_ue_ in anatomical (*Left*) and network space (*Middle*). *Right:* Distribution of top 5% FC increases across all networks.

### Plasticity Was Restricted to the Somatomotor and Cingulo-opercular Networks

The analyses presented thus far focus on changes in connectivity between L-SM1_ue_ and the remainder of the cerebral cortex. To examine whole-brain patterns of FC changes induced by casting, we extracted rs-fMRI signals from individual-specific cortical, subcortical and cerebellar parcels (Cast1: 744 parcels, Cast2: 733; Cast3: 761) and examined FC changes in all possible pairwise connections (Fig. S3). We displayed the 50 largest changes in FC (25 greatest increases and 25 greatest decreases) in anatomical space (Fig. 3*A*, S3). Disuse-driven FC changes involved L-SM1_ue_ more than expected by chance (Cast1: 9/50 connections, P = 0.018; Cast2: 37/50, P < 0.001; Cast3: 39/50, P < 0.001). Figure 3*B* shows the total magnitude of whole-brain FC change for each vertex/voxel. The total magnitude of whole-brain FC change was significantly greater in L-SM1_ue_ than in the remainder of the brain (Fig. 3*B*, S3; Cast1: P = 0.021; Cast2: P < 0.001; Cast3: P = 0.005).

**Fig. 3.**
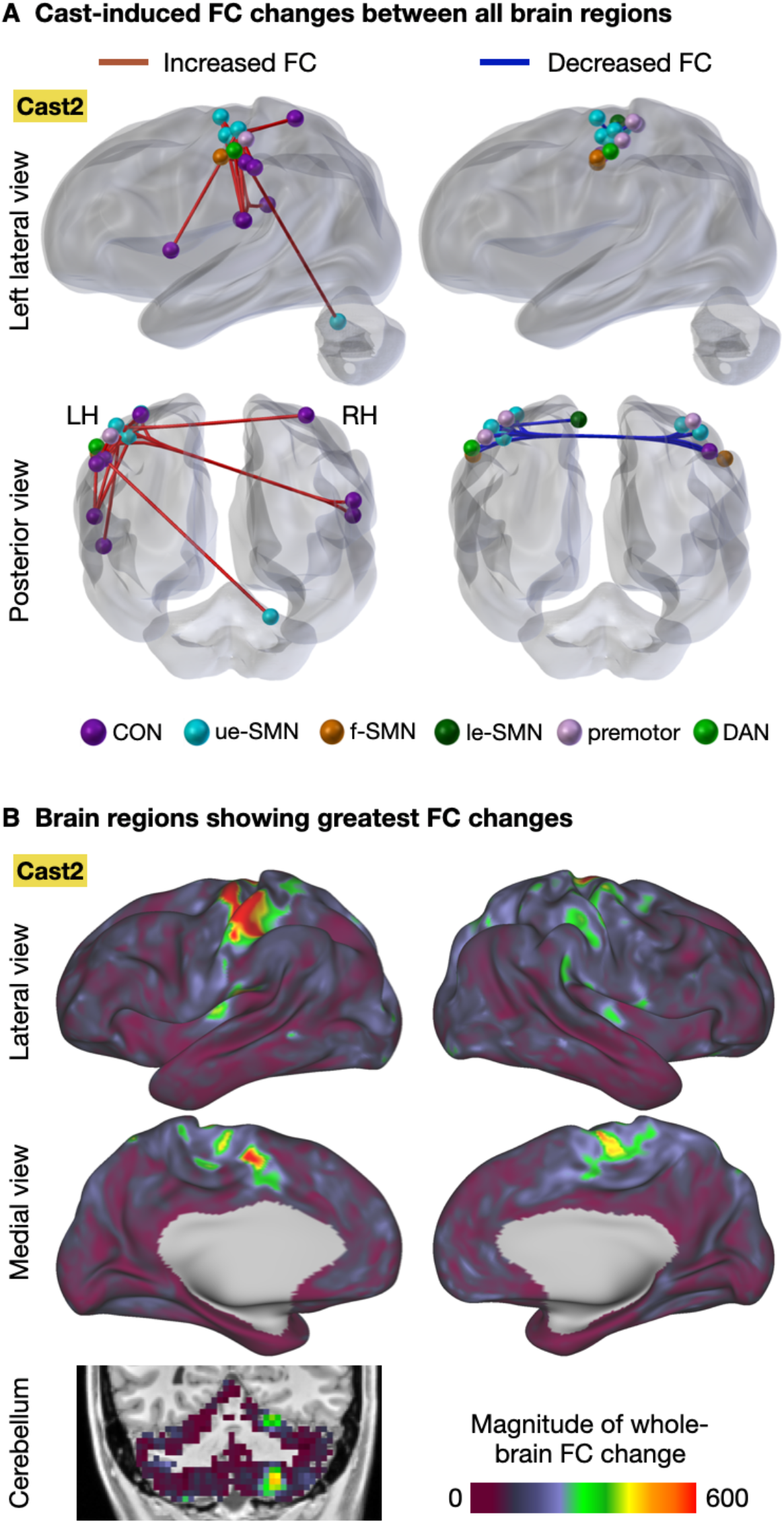
Whole-brain Analyses of Disuse-driven Functional Connectivity (FC) Changes. (A) Changes in functional connectivity (FC) were computed for all pairs of cortical, subcortical and cerebellar parcels (mean Cast FC – mean Pre FC). The 50 most altered functional connections (top 25 increases, top 25 decreases) are shown for an example participant (Cast2; all participants, Fig. S3). (B) For each vertex/voxel, the magnitude of whole-brain FC change was computed as the sum of squared FC changes between that vertex and every other gray-matter vertex/voxel. Shown for an example participant (Cast2; all participants shown in Fig. S4).

### Increased Connectivity with the CON Depended on a Recent History of Disuse

To distinguish between increases in connectivity due to wearing a cast during MRI scans (i.e., an altered behavioral state) and increases in connectivity due to a history of having been casted (i.e., disuse-driven plasticity), we conducted a control experiment in which participants wore casts during 30-minute scans but were not casted during their daily lives. In both cases, we compared FC between L-SM1_ue_ and the left dorsal anterior cingulate cortex (L-dACC; Fig. S4*A*), a key node of the CON, in cast and no-cast conditions. As suggested by the whole-brain network analyses (Fig. 2*B*), FC between L-SM1_ue_ and L-dACC was significantly increased during the cast period (Fig. S4*B*). When participants wore casts during scans, but were not casted during their daily lives, no changes in FC were observed (Fig. S4*C*). This control experiment indicated that increased FC between the disused motor circuitry and the CON depended on participants’ recent history of disuse, not their current state of casting.

### Network-Specific Connectivity Changes Reflect Spontaneous Activity Pulses

Increases in FC with L-SM1_ue_ showed a similar anatomical distribution to the spontaneous activity pulses we described previously (31) (Fig. S5*A-B*). Additionally, FC between L-SM1_ue_ and the CON was highly correlated with the number of pulses detected in L-SM1_ue_ during each scan (Fig. S5*C*; Cast1: r=0.74, Cast2: r=0.57, Cast3: r=0.73; all participants: P<0.001). These observations led us to suspect that FC between L-SM1_ue_ and the CON was increased during the cast period because both regions showed synchronized spontaneous activity pulses.

To test if spontaneous activity pulses could explain observed increases in FC, we implemented a pulse censoring strategy, in which frames surrounding each detected pulse (13.2 s before – 17.6 s after each pulse peak) were excluded from FC calculations (Fig. 4*A*). This approach is similar to the censoring strategy commonly used to correct the effects of head-motion on FC (32). Censoring pulses partially reduced the FC changes observed during casting (Fig. 4*B*; Cast1 −55%, P = 0.14; Cast2: −28%, P = 0.02; Cast3: −21%, P = 0.04; one-tailed t-test). Additionally, the spatial pattern of FC changes caused by pulse censoring (Censored – Uncensored; Fig. 4*C*, S6) was negatively correlated with FC changes during casting (Cast1: r = −0.35, Cast2: r = −0.90, Cast3: r = −33; all participants: P < 0.001).

**Fig. 4.**
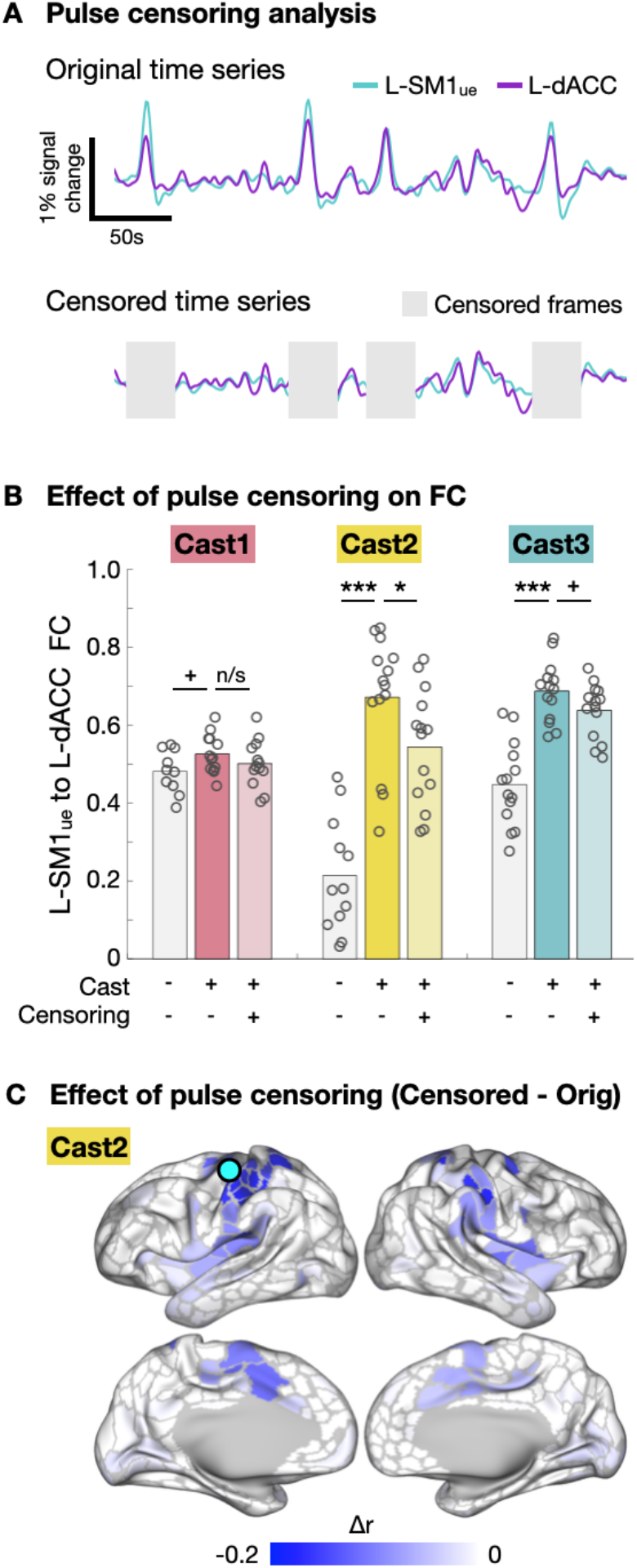
Censoring Pulses from Functional Connectivity (FC) Measurements. (A) Schematic explaining censoring approach used to remove detected pulses from FC calculation. (B) Censoring pulses partially reduced the difference between Pre and Cast scans. ^+^P < 0.1, *P < 0.05, ***P < 0.001. (C) Subtraction map showing reductions in L-SM1_ue_ FC due to pulse censoring. The spatial pattern of FC changes due to pulse censoring was significantly negatively correlated with FC changes during casting (Cast1: r = −0.35, Cast2: r = −0.90, Cast3: r = −0.33; all participants: P < 0.001)

Censoring pulses did not entirely eliminate the effect of casting on FC. The residual effect of casting after pulse censoring was negatively spatially correlated with the effects of pulse censoring (Fig. S6; Cast1: spatial correlation, r = −0.23, Cast2: r = −0.76, Cast3: r = −0.18; all participants: P < 0.001), suggesting that the residual effect may have been driven by additional pulses that were too small to distinguish from ongoing spontaneous activity fluctuation. Pulses showed a wide distribution of magnitudes (Fig. 5*A*; range: 0.4 – 1.2% rs-fMRI signal change). Because pulses were detected using a threshold-based approach (31), pulses smaller than 0.4% fMRI signal change could not be detected and censored. To test if smaller pulses that escaped detection could explain the residual increased FC between L-SM1_ue_ and L-dACC after pulse censoring, we simulated a full distribution of pulse magnitudes by adding the mean pulse time series from L-SM1_ue_ and L-dACC to the baseline rs-fMRI signals recorded from these regions prior to casting (Fig. 5*B*). Simulated pulse magnitudes were drawn from a log-normal distribution, fit to the magnitude distribution of detected pulses using a least-squares approach (Fig. 5*A*; see S7 for triangular and exponential distributions). Adding simulated pulses caused FC increases with a similar effect size (Cast1: Δr = +0.07, P = 0.013; Cast2: +0.40, P < 0.001; Cast3: +0.19, P < 0.001) to the FC increases observed during casting (Fig. 5*C*). Applying the same pulse censoring strategy described above to the simulated pulses partially reduced the increases in FC (Fig. 5*C*; Cast1: −79%, Cast2: −58%, Cast3: −69%; all participants: P < 0.001), similar to the effect of pulse censoring on the actual data (Fig. 4*B*). Adding simulated pulses to all brain regions, using mean pulse time series specific to each region, recreated the anatomical distribution of FC changes observed during casting (Fig. 5*D*, S8; Cast1: r = 0.44, Cast2: r = 0.95, Cast3: r = 0.64; all participants: P < 0.001). Thus, pulse with a full distribution of magnitudes could account for observed FC changes during casting and the partial reversal of FC changes after pulse censoring.

**Fig. 5.**
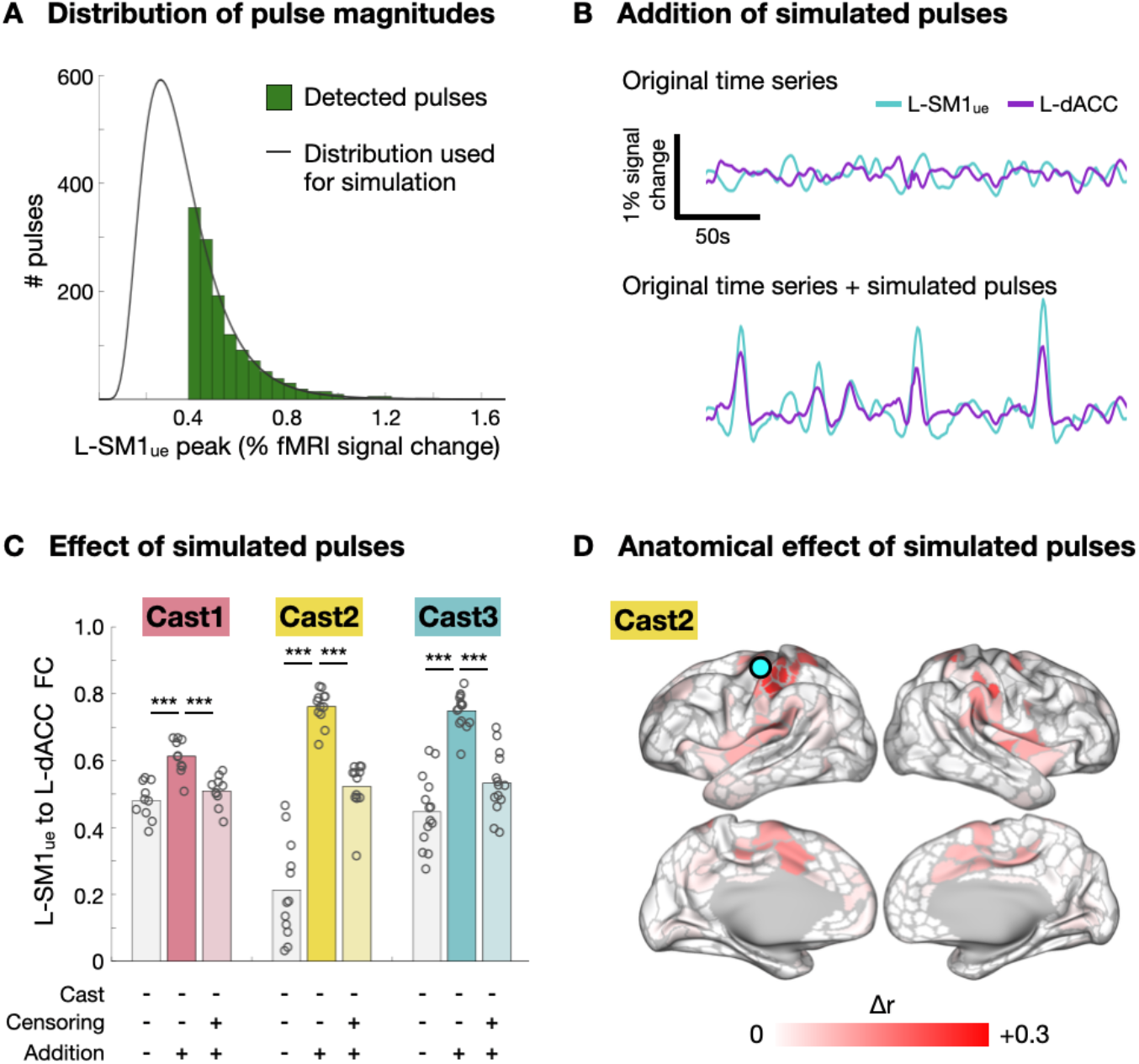
Addition of Simulated Pulses. (A) Histogram of pulse magnitudes (peak fMRI signal), pooled across participants. A log-normal distribution (black line) was fit to the data using a least-squares approach. (B) *Top:* Example resting-state functional MRI (rs-fMRI) signals from disused motor cortex (L-SM1_ue_) and the dorsal anterior cingulate cortex (L-dACC) recorded prior to casting (Pre). *Bottom:* Same time series with simulated pulses added at random time points. Simulated pulse magnitudes were drawn from the full distribution shown in panel A. (C) Adding simulated pulses increased FC between L-SM1_ue_ and L-dACC (***P<0.001). Censoring simulated pulses partially reduced FC increases (***P<0.001). (D) Subtraction map showing increases in L-SM1_ue_ FC caused by simulated pulses in an example participant (Cast2; all participants, Fig. S7).

We also utilized the pulse censoring and pulse addition analyses to test if pulses could explain observed decreases in FC within the somatomotor system. Pulse censoring did not significantly reverse the decrease in FC between L-SM1_ue_ and R-SM1_ue_ observed during the cast period (Fig. S9). Adding simulated pulses to baseline time series decreased FC between L-SM1_ue_ and R-SM1_ue_, but this effect was much smaller than that observed during casting (Fig. S9).

## Discussion

### Disuse Drives Plasticity of Functional Networks

Daily 30-minute scans of rs-fMRI before, during, and after casting revealed that disuse not only causes plasticity within the primary motor cortex (4, 31), but also increased functional connectivity between disused circuits and executive control regions in the CON. These increases in FC would have been difficult to observe using the focal recording techniques traditionally employed to study plasticity. Thus, understanding the total impact of disuse on brain function requires consideration of plasticity at multiple spatial scales.

The high degree of network specificity seen in disuse-driven plasticity is a powerful demonstration of the validity of individual-specific rs-fMRI network parcellations. It could have been the case that casting produced a complex pattern of FC changes involving many brain systems. Instead, we found that virtually all of the regions showing increased connectivity with the disused motor cortex belonged to a single network, the CON. Thus, large-scale functional networks seem to be a fundamental unit of brain organization, important not only for understanding patterns of activation during cognitive processes, but also for understanding whole-brain patterns of plasticity.

### Spontaneous Activity Pulses Alter Functional Connectivity

The network-specific increases in FC between the disused L-SM1_ue_ and the CON likely reflect the emergence of synchronized spontaneous activity pulses in these regions during casting. FC is a measurement of the temporal correlation of spontaneous activity between brain regions. Two complementary analyses (pulse censoring and pulse addition) suggested that pulses contributed to increases in measured FC between disused motor regions and the CON. Many studies have reported increased FC in neuropsychiatric conditions, e.g., Parkinson’s disease (33) and Major Depressive Disorder (34). Perhaps some previously reported FC increases could reflect spontaneous activity pulses, similar to those caused by limb disuse. Going forward, spontaneous activity pulses, and the network-specific FC changes they produce, could serve as a biCast3ker of recent disuse and states of heightened plasticity (35).

Decreases in connectivity within the somatomotor system were not fully explained by spontaneous activity pulses. Although the pulse addition analyses showed that spontaneous activity pulses could contribute to decreases in connectivity observed during casting, both the pulse censoring and pulse addition analyses suggested that at least part of the observed decrease in connectivity occurred independently of pulses. Loss of typical co-activation of L-SM1_ue_ and R-SM1_ue_ during daily behavior (31) likely contributed to cast-driven connectivity decreases, consistent with the widely held hypothesis that FC is maintained by a Hebbian-like mechanism that depends on co-activation of brain regions during behavior (19, 36–39).

### Disuse Impacts Cingulo-opercular Network (CON) Important for Executive Control

The importance of control systems in the human brain has been recognized for several decades (7), but our understanding of these systems has changed dramatically over time and continues to evolve (40). Disentangling the roles of two networks mediating executive control, the cingulo-opercular and frontoparietal networks, has relied largely on complex task designs such as task-switching (21) and slow-reveal paradigms (41). Here, we have demonstrated that a new approach, whole-brain precision functional mapping combined with anatomically specific disuse paradigms, can shed new light on the network interactions that support human cognition and behavior. Although the brain contains several suspected control networks beyond the CON, such as the frontoparietal (FPN), dorsal attention (DAN), ventral attention (VAN) and salience (SAL) networks, disused motor regions showed increased FC only with the CON. The CON is thought to initiate goal-directed behaviors and maintain executive control settings in relation to task objectives (1, 19). The increased FC between disused L-SM1_ue_ and the CON during casting suggests either 1) circuits within the CON were also disused during casting, 2) the CON helps to maintain disused motor circuitry, or 3) spontaneous activity pulses can spread along functional connections to neighboring brain regions in network space. Spontaneous activity pulses may have emerged in the CON because its use was also reduced during casting. We previously suggested that spontaneous activity pulses found in motor regions may have been caused by disuse-driven changes in local inhibitory physiology (31, 42). The CON is generally thought to represent task sets, abstract parameters and motor programs governing goal-driven behaviors (1, 19). Task sets can be applied to multiple motor effectors (e.g., right hand, left index finger, etc.), so it would be reasonable to think that casting one extremity would not cause disuse of the CON. However, at least two sets of circuits within the CON may have been disused during casting: circuits that represent bimanual behaviors and circuits that convey task sets to effector-specific downstream regions (e.g., L-SM1_ue_). During casting, many of the task sets previously used to control the dominant upper extremity (e.g., inserting a key into a lock) could have been used to control the non-dominant extremity. However, task sets requiring bimanual coordination (e.g., fastening a belt) could not be applied to an alternative set of motor effectors and may have become disused. Another set of CON circuits that may have been disused are the circuits that convey task set information to effector-specific brain regions. The CON contains both somatotopically organized regions (e.g., SMA, SII) and non-somatotopic regions (e.g., anterior insula). Recent work shows that the SMA and SII, as well as regions of the basal ganglia, may act as hubs that connect the rest of the CON to the somatomotor network (27, 43). Such circuits could potentially undergo disuse during casting of one extremity.

An alternative possibility is that inputs from the CON to L-SM1_ue_ served a homeostatic function during disuse, helping to maintain motor circuits that are typically maintained through active use. We previously suggested that spontaneous activity pulses may help to maintain the organization of disused brain circuits (31). Perhaps pulses are triggered by the CON, which is the system typically responsible for initiating activation of the somatomotor network (1).

Spontaneous activity pulses may also have originated in the disused somatomotor circuits and spread along functional connections to brain regions that were not affected by casting. The CON is immediately adjacent to the somatomotor network in network space (Fig. 2*A*). Previous work in mice found that spontaneous activity bursts in somatosensory cortex, caused by local pharmacogenetic suppression of inhibitory interneurons, spread to other functionally connected regions not targeted by the experimental manipulation (44). An interesting question for future research is whether the CON would show spontaneous activity pulses following disuse of brain systems other than the somatomotor network, such as networks supporting visual and auditory processing, spatial navigation, or social cognition.

### Two Complementary Mechanisms for Functional Network Plasticity

Extensive whole-brain imaging before, during and after casting revealed that disuse-driven spontaneous activity pulses occur not only in primary motor and somatosensory areas, but also in higher-order brain regions responsible for executive control over behavior. The emergence of spontaneous activity pulses during casting produced increases in FC that were highly focal in network space, specifically occurring between disused motor regions and the CON. Decreases in connectivity between disused motor circuits and the remainder of the somatomotor system, however, were not explained by spontaneous activity pulses. Thus, disuse may drive network plasticity through two complementary mechanisms. Decreased coactivation of brain regions during disuse might drive Hebbian-like disconnection between disused and still-active somatomotor circuits. Spontaneous activity pulses, which contributed to increased connectivity with the CON, may help preserve disused circuits for future reintegration and use. Together, these two network plasticity mechanisms—Hebbian-like and pulse-mediated—may represent a network-focal “stand-by mode” that allows the brain to isolate disused circuits while simultaneously protecting them from premature functional degradation.

## Acknowledgements

We thank Kristen M. Scheidter and Annie L. Nguyen for help in collecting MRI data and Ryan V. Raut for his comments on the manuscript. This work was supported by NIH grants NS110332 (D.J.N.), NS088590 (N.U.F.D.), TR000448 (N.U.F.D.), MH96773 (N.U.F.D.), MH122066 (N.U.F.D.), MH1000872 (T.O.L.), MH112473 (T.O.L.), NS090978 (B.P.K.), MH104592 (D.J.G.), NS080675 (M.E.R.), NS098577 (to the Neuroimaging Informatics and Analysis Center); the US Department of Veterans Affairs Clinical Sciences Research and Development Service grant 1IK2CX001680 (E.M.G.); Kiwanis Neuroscience Research Foundation (N.U.F.D.); the Jacobs Foundation grant 2016121703 (N.U.F.D.); the Child Neurology Foundation (N.U.F.D.); the McDonnell Center for Systems Neuroscience (D.J.N., T.O.L., A.Z., B.L.S., and N.U.F.D.); the McDonnell Foundation (S.E.P.), the Mallinckrodt Institute of Radiology grant 14-011 (N.U.F.D.); the Hope Center for Neurological Disorders (N.U.F.D., B.L.S., and S.E.P.).

## Author Contributions

D.J.N., T.O.L., C.R.H., A.Z.S., and N.U.F.D. designed the study. D.J.N., C.R.H., J.M.H., M.O., and N.U.F.D. collected the data. D.J.N. and E.M.G. analyzed the data. D.J.N., E.M.G., T.O.L., N.A.S., D.F.M., S.J.G., A.Z., A.N.N., B.A., A.R.C., B.P.K., D.J.G., M.E.R., S.E.P., A.Z.S., and N.U.F.D. interpreted the results. D.J.N. and N.U.F.D. wrote the manuscript with input from all other authors.

## METHODS

### Human Participants

Participants were three healthy, adult volunteers. Details are described in Newbold *et al*. (31).

### Experimental Intervention

We constrained the dominant upper extremity for two weeks by fitting each participant with a fiberglass cast. Casts extended from just below the shoulder to past the fingertips. Details are described in Newbold *et al*. (31).

### MRI Acquisition

Participants were scanned every day of the experiment for 42-64 consecutive days. We collected 30 minutes of resting-state functional MRI (rs-fMRI) data every day of the experiment, as well as also task fMRI and T1- and T2-weighted structural scans. Acquisition parameters and procedures are detailed in Newbold *et al*. (31).

### MR Image Processing

Preprocessing of structural and functional images, denoising of resting-state fMRI data, and cortical surface projection were performed previously (31). Fully processed data are available in the Derivatives folder of the Cast-Induced Plasticity dataset on OpenNeuro (www.openneuro.org/datasets/ds002766).

### ROI Selection

The region of the left primary somatomotor cortex controlling the casted upper extremity (L-SM1_ue_) was located individually in each participant using task fMRI. Details of the task analysis and region of interest (ROI) selection are described in Newbold *et al*. (31). The L-SM1_ue_ ROI was used here to generate seed maps (Fig. 1, S1), to analyze spatial specificity of whole-brain functional connectivity changes (Fig. 3*B*, S3) and to measure FC between L-SM1_ue_ and the dorsal anterior cingulate cortex (dACC) for the removable cast control experiment (Fig. S4), the pulse censoring analysis (Fig. 4) and the pulse addition analysis (Fig. 5, S7).

Individual-specific sets of parcels spanning the entire cerebral cortex were previously generated for each participant using a functional connectivity gradient-based approach, and each parcel was assigned to one of 17 canonical functional networks using a graph theory-based community detection algorithm (22, 31). Cortical parcels were used here to display functional connectivity changes in anatomical (Fig. 1–2, 5, S1–S2, S4–S6) and network space (Fig. 2, S2). We also generated a set of subcortical and cerebellar parcels by assigning each voxel to the functional network with which it showed the strongest functional connectivity and then grouping adjacent voxels with matching network assignments into parcels. Sets of cortical, subcortical and cerebellar parcels were used to examine changes in FC between all pairs of brain regions (Fig. 3, S2).

A subset of parcels corresponding to the left primary somatomotor cortex (L-SM1_ue_) was previously generated by selecting all left-hemisphere parcels assigned to the upper-extremity somatomotor network, excluding any parcels that fell outside of the pre- and post-central gyri (31). Functional connections between L-SM1_ue_ parcels and all other parcels were averaged to generate parcel-wise seed maps (Fig. 1–2, 5, S1–S2, S4–S6) and to quantify the spatial specificity of FC changes between all cortical, subcortical and cerebellar parcels (Fig. 3, S3).

Individual-specific ROIs in the left dorsal anterior cingulate cortex (L-dACC) were selected using pulse analysis of variance (ANOVA) maps, which were previously generated for each participant (31). ROIs were selected within a large anatomical region, automatically labeled by FreeSurfer (union of caudal anterior cingulate gyrus, posterior cingulate gyrus, and superior frontal gyrus). We first located the vertex showing the maximum ANOVA F-statistic and then iteratively expanded the ROI by selecting neighboring vertices in descending order of F-statistic, until the ROI included 100 vertices. The L-dACC ROI was used to measure FC between L-SM1_ue_ and the L-dACC for pulse censoring (Fig. 4) and pulse addition analyses (Fig. 5, S7).

### Functional Connectivity Measurement

Mean rs-fMRI time series were extracted from the L-SM1_ue_ ROI and each cortical, subcortical and cerebellar parcel by averaging the time series of all vertices/voxels within the ROI/parcel. Vertex-wise functional connectivity (FC) seed maps were generated for each session by computing a Pearson correlation between the mean rs-fMRI time series from the L-SM1_ue_ ROI and the time series from every vertex. Frames with high head motion, identified previously (31), were excluded from FC calculations. Vertex-wise seed maps were averaged across sessions before, during and after casting (Pre, Cast, Post; Fig. 1, S1).

FC between all pairs of cortical, subcortical and cerebellar parcels was measured as pairwise correlations of all mean rs-fMRI time series, excluding high-motion frames, producing a correlation matrix for each session. Correlation matrices were averaged over Pre, Cast and Post period. FC changes during casting and recovery were computed as the differences between correlation matrices (casting = Cast – Pre; recovery = Post – Cast). Full Cast – Pre differences matrices are shown in Figure S3. Parcel-wise difference seed maps showing changes in L-SM1_ue_ FC during casting and recovery (Fig. 1–2, S1–S2) were computed by averaging together columns of the difference matrices corresponding to the L-SM1_ue_ parcels (see *ROI selection*, above).

### Spring Embedding

Cortical parcels were displayed in network space using force-directed (“spring-embedded”) graphs(45), generated with Gephi (https://gephi.org/). Graph weights were taken from parcel-wise correlation matrices averaged across all sessions prior to casting (Pre). Graphs were thresholded to include only the top 0.2% of pairwise functional connections. We initially examined graphs using an edge threshold of 0.1%, but several parcels with large FC changes were disconnected from the main graph. Thus, we selected the 0.2% connection threshold to display all key findings (Fig. 2, S2).

### Functional Connectivity Changes by Network

To test which functional networks showed large changes in FC with L-SM1_ue_ during casting, parcel-wise L-SM1_ue_ seed maps were compared to individual-specific network maps. Parcel-wise L-SM1_ue_ difference seed maps (Cast – Pre) were thresholded to contain the top 5% of parcels showing FC increases/decreases. FC increases (Cast > Pre) and decreases (Cast < Pre) were examined separately. The number of supra-threshold parcels in each cortical network is shown in Figure 2. Alternative thresholds (top 1%, 10% and 20%) all yielded similar results. Networks containing a greater number of FC increases/decreases than expected by chance were identified by comparing the number of supra-threshold parcels found using each participant’s true network map to the number of parcels found using spatially permuted network maps (see *Statistical analyses*, below).

### Whole-brain Analyses of Functional Connectivity Changes

The spatial specificity of whole-brain plasticity was examined by comparing changes in FC between all cortical, subcortical and cerebellar parcels to individual-specific functional network maps. We examined the top 50 changes in FC (25 greatest increases and 25 greatest decreases). A change in FC between two parcels was counted as involving L-SM1_ue_ if either of the two parcels was an L-SM1_ue_ parcel. The number L-SM1_ue_ changes found using each participant’s true network map was compared to the number of L-SM1_ue_ changes found using spatially permuted network maps (see *Statistical analyses*, below).

To examine whole-brain FC changes at a finer spatial resolution, we examined differences in FC between all cortical, subcortical and cerebellar vertices/voxels. We computed the magnitude of whole-brain FC change between each vertex/voxel as the sum of squared FC changes between that vertex/voxel and every other gray-matter vertex/voxel (Fig. 3, S3). We tested if whole-brain FC change was higher in L-SM1_ue_ than in the rest of the brain by comparing the mean magnitude of whole-brain FC change inside of each participant’s true L-SM1_ue_ ROI to the mean magnitude of whole-brain FC change inside of spatially rotated ROIs (see *Statistical analyses*, below).

### Spatial and Temporal Comparisons of Pulses and FC Changes

To generate a parcel-wise map of spontaneous activity pulses, we extracted rs-fMRI time series from every cortical, subcortical and cerebellar parcel surrounding each detected pulse (13.2 seconds before to 17.6 seconds after each pulse peak). Pulse peaks were detected previously (31). We then performed an ANOVA on the extracted time series (Fig. S5*B*). The spatial distribution of ANOVA F-statistics was compared to the spatial distribution of L-SM1_ue_ FC changes using a Pearson correlation across parcels.

We also compared the number of pulses detected during each rs-fMRI scan (31) to the FC measured between L-SM1_ue_ and L-dACC (Fig. S4*C*), using a Pearson correlation.

### Pulse Censoring Analyses

To assess the contribution of large, detectable spontaneous activity pulses (>0.4% rs-fMRI signal change) to FC changes observed during casting, frames surrounding each pulse (13.2 seconds before to 17.6 seconds after each pulse peak) were excluded from FC calculations. Pulse detection criteria were identical to those used previously (31). We plotted FC measured between L-SM1_ue_ and L-dACC during each session before casting (Pre), during casting (Cast) and during casting with pulses excluded (Cens; Fig. 4*B*). We compared Cast vs. Pre FC measurements and Cens vs. Cast FC measurements using t-tests (see *Statistical analyses*, below). We also generated difference seed maps showing FC changes between L-SM1_ue_ and all cortical parcels due to censoring (Cens – Cast; Fig. 4*C*, S6).

### Pulse Addition Analyses

To assess the contribution to FC changes of pulses with a hypothetical full magnitude distribution, including pulses that were too small (<0.4% rs-fMRI signal change) to distinguish from ongoing spontaneous activity fluctuations, we created simulated rs-fMRI time series containing a full distribution of pulses. We first generated a histogram showing the magnitude distribution of detected pulses. Magnitudes were grouped into bins with a width of 0.5%-signal-change. We then fit a log-normal distribution to the observed pulse magnitudes using a least-squares approach (Fig. 5*A*). This provided an estimate of the full pulse magnitude distribution, including pulses that were too small to detect. We then added the mean pulse time series to the baseline (Pre) rs-fMRI time series, drawing pulse magnitudes from the full log-normal distribution (example shown in Fig. 5*B*). The number of pulses added was selected so that the number of large pulses (>0.4% signal change) would match the number of pulses detected in each participant in an average Cast scan. We then computed FC of the baseline time series (Pre) and the time series with added pulses (Sim; Fig. 5*C*). Finally, we applied the same pulse censoring strategy described above to the simulated time series and computed FC after censoring (Cens; Fig. 5*C*). Because pulse magnitudes were drawn from a full log-normal distribution, only a portion of the added pulses were detected and censored. We compared Sim vs. Pre FC measurements and Cens vs. Sim FC measurements using t-tests (see *Statistical analyses*, below). The simulation and censoring procedures were repeated using triangular and exponential magnitude distributions (Fig. S7). We also generated difference seed maps showing FC changes between L-SM1_ue_ and all cortical parcels due to pulse simulation (Sim – Pre; Fig. 5*D*, S8).

### Statistical Analyses

All statistical tests were performed identically for each participant. Whenever appropriate, we used simple parametric statistical tests:

- Spatial similarity between anatomical maps was tested using a Pearson correlation across parcels (Cast1: d.f. = 566; Cast2: 578; Cast3: 624). Spatial correlations were computed between the following maps:

○ FC changes during casting (Cast – Pre) vs. FC changes during recovery (Post – Cast; Fig. 1*B*)
○ FC changes during casting (Cast – Pre) vs. pulse ANOVA (F-statistic; Fig. S5)
○ FC changes during casting (Cast – Pre) vs. FC changes due to pulse censoring (Cens – Cast; Fig. 4*C*, S6)
○ FC changes due to pulse censoring (Cens – Cast) vs. residual FC changes after pulse censoring (Cens – Pre; Fig. S6)
○ FC changes during casting (Cast – Pre) vs. FC changes due to simulated pulses (Sim – Pre; Fig. 5*D*, S8)
- The number of pulses detected during each scan was compared to the FC measured between L-SM1_ue_ and L-dACC using a Pearson correlation (Cast1: d.f. = 45; Cast2: 61; Cast3: 40; Fig. S5*C*).
- FC between L-SM1_ue_ and L-dACC measured during each session of the cast period (Cast1: n = 13 sessions; Cast2: 13; Cast3: 14) was compared to FC during each session of the pre period (Cast1: n = 10; Cast2: 12; Cast3: 14) using a two-sided, unpaired t-test (Cast1: d.f. = 21, Cast2: 23, Cast3: 26; Fig. S5*B*).
- FC between L-SM1_ue_ and L-dACC measured using the full time series (motion-censored only) of each session of the cast period (Cast1: n = 13 sessions; Cast2: 13; Cast3: 14) was compared to FC measurements of the same sessions excluding pulses (motion- and pulse-censored) using a one-sided, paired t-test (Cast1: d.f. = 24, Cast2: 24, Cast3: 26; Fig. 4*B*).
- FC between L-SM1_ue_ and L-dACC measured during each session of the pre period (Pre; Cast1: n = 10; Cast2: 12; Cast3: 14) was compared to FC measurements of the same sessions after adding simulated pulses (Sim) using a two-sided, paired t-test (Cast1: d.f. = 18, Cast2: 22, Cast3: 26; Fig. 5*C*).
- FC between L-SM1_ue_ and L-dACC measured using the full time simulated series (Sim; motion-censored only; Cast1: n = 10; Cast2: 12; Cast3: 14) was compared to FC measurements of the same sessions excluding pulses (motion- and pulse-censored) using a two-sided, paired t-test (Cast1: d.f. = 18, Cast2: 22, Cast3: 26; Fig. 5*C*).

When parametric statistical tests were not appropriate to test a specific hypothesis, we tested results against a null distribution generated via permutation resampling. In each case, our null hypothesis was that observed effects had no spatial relationship to ROIs/functional networks and any overlap occurred by chance. For vertex-wise ROIs, we modeled the null hypothesis by rotating the ROI around the cortical surface 1,000 times (46, 47). For functional networks, we modeled the null hypothesis by permuting the network assignments of parcels 1,000 times. Each permuted ROI/network map was used exactly as the actual map in order to compute a null distribution for the value of interest. The P-value reported for each test represents the two-sided probability that a value in the null distribution has a greater magnitude than the observed value. Permutation resampling was used to generate null distributions for the following values:

- Overlap of each functional network with increases in L-SM1_ue_ FC during casting (Cast > Pre; number of parcels in top 5%; Fig. 2*B*)
- Overlap of each functional network with decreases in L-SM1_ue_ FC during casting (Cast > Pre; number of parcels in top 5%; Fig. 2*C*)
- Overlap of L-SM1_ue_ parcels with the 50 largest changes in FC between all pairs of cortical, subcortical and cerebellar parcels (Fig. 3*A*, Fig. S3)
- Magnitude of whole-brain FC change in the L-SM1_ue_ ROI (Fig. 3*B*, S3)

Since we tested the overlap of L-SM1_ue_ FC increases/decreases with 17 different functional networks, a Benjamini-Hochberg procedure was applied to correct for multiple comparisons, maintaining false discovery rates < 0.05. Each of the three participants constituted a separate replication of the experiment, rather than multiple comparisons, so no correction was necessary for tests repeated in each participant.

### Data Visualization

Regions of interest and whole-brain pulse maps were shown on cortical surfaces generated by FreeSurfer (48) and Human Connectome Project (HCP) Workbench software packages (49). These images were rendered using HCP Workbench (49). Figures showing the greatest differences in functional connectivity across the entire brain in inflated anatomical space (Fig. 3, S3) were rendered using Blender, a free and open-source 3D modeling software package (www.blender.org). To make the large number of changed connections included in these images visually compact, connections following similar spatial trajectories were drawn toward each other using a previously published mean-shift edge bundling algorithm (50). All other figures were produced using Matlab (www.mathworks.com).

### Data Availability

This study used our previously published dataset (31), available on the OpenNeuro database (www.openneuro.org/datasets/ds002766).

### Code Availability

All code needed to reproduce our analyses is available on Gitlab (https://gitlab.com/DosenbachGreene/cast-whole-brain).

---

**Fig. S1.**
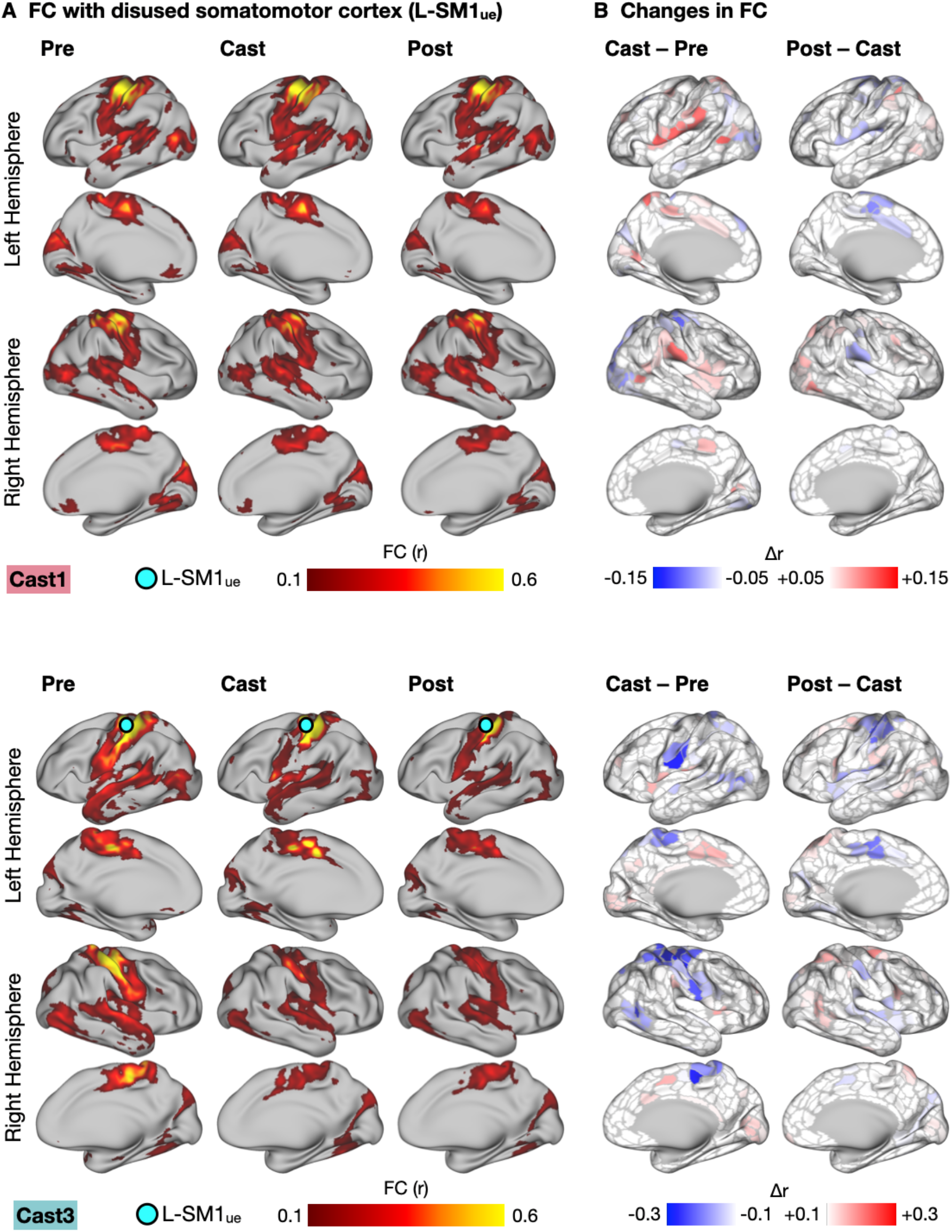
Functional Connectivity (FC) Changes Involving the Disused Somatomotor Cortex (L-SM1_ue_) in Anatomical Space (All Participants). (A) Seed maps showing FC between L-SM1_ue_ and the remainder of the cerebral cortex in Cast1 (top) and Cast3 (bottom). Seed maps were averaged across sessions before, during and after casting (Pre, Cast, Post). (B) Subtraction maps showing changes in FC during casting (Cast – Pre) and recovery (Post – Cast). Differences are shown on sets of 506 (Cast1) and 626 (Cast3) individual-specific cortical parcels. For Cast1 and Cast2, these parcels were previously generated and published (Gordon, 2017). We applied identical methods to generate a set of parcels for Cast3.

**Fig. S2.**
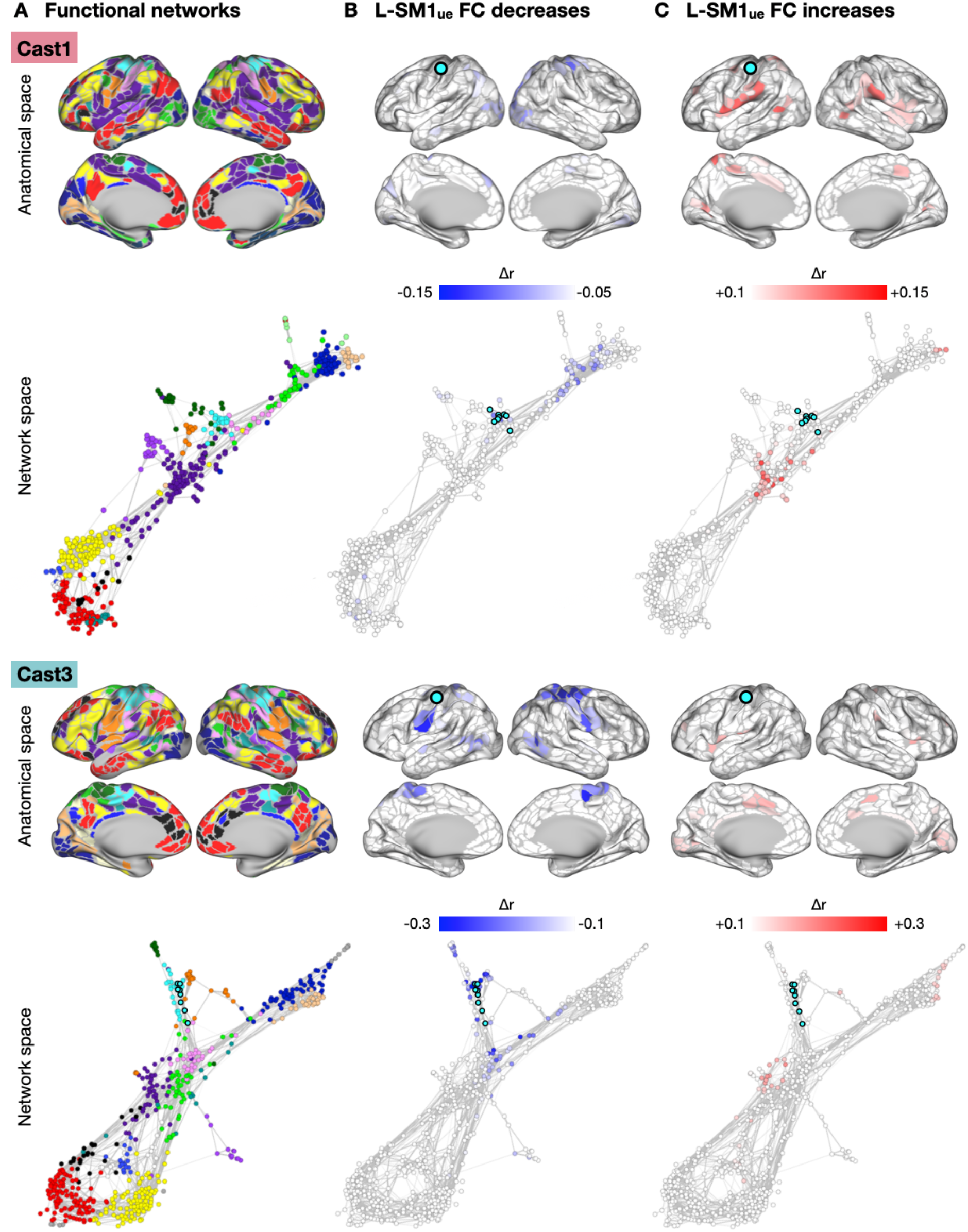
Functional Connectivity (FC) Changes Involving the Disused Somatomotor Cortex (L-SM1_ue_) in Anatomical and Network Space in Cast1 and Cast3. (A) Maps of 17 canonical functional networks in Cast1 (top) and Cast3 (bottom). Networks are shown in anatomical space (inflated cortical surfaces) and network space (spring-embedded graphs based on Pre scans). See Figure 2 for network color key. (B) Subtraction maps showing disuse-driven decreases in functional connectivity (FC) with left somatomotor cortex (L-SM1_ue_) in anatomical space and network space. (C) Subtraction maps showing disuse-driven increases in FC with L-SM1_ue_ in anatomical and network space.

**Fig. S3.**
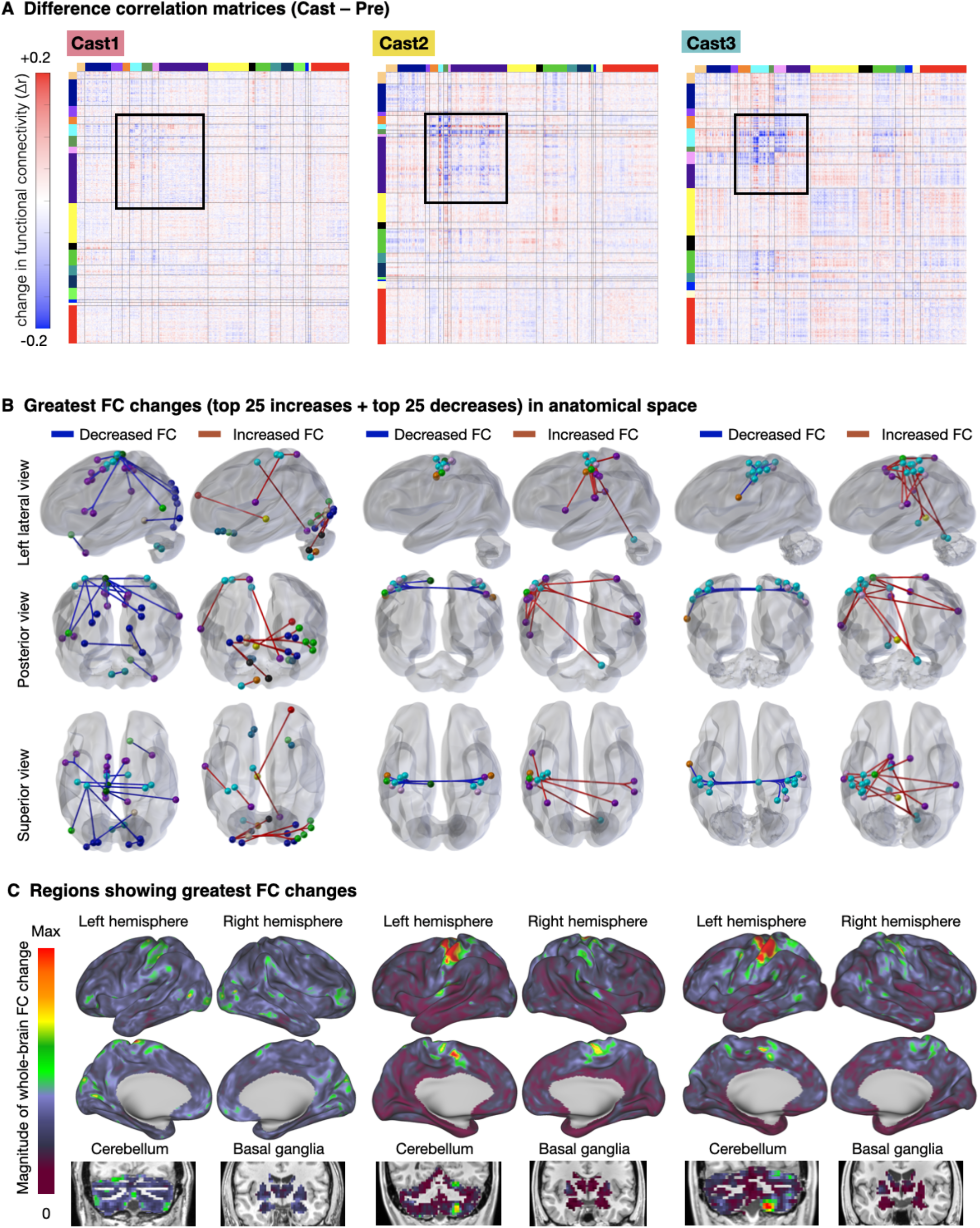
Whole-brain Patterns of Disuse-driven Plasticity. (A) Matrices showing changes in functional connectivity (FC) between all pairs of individual specific cortical, subcortical and cerebellar parcels during casting (Cast1: 744 parcels, Cast2: 733; Cast3: 761). (B) 50 largest changes in FC shown in anatomical space. (C) Magnitude of whole-brain FC change, computed as the sum of squared FC changes between each vertex/voxel and every other gray-matter vertex/voxel.

**Fig. S4.**
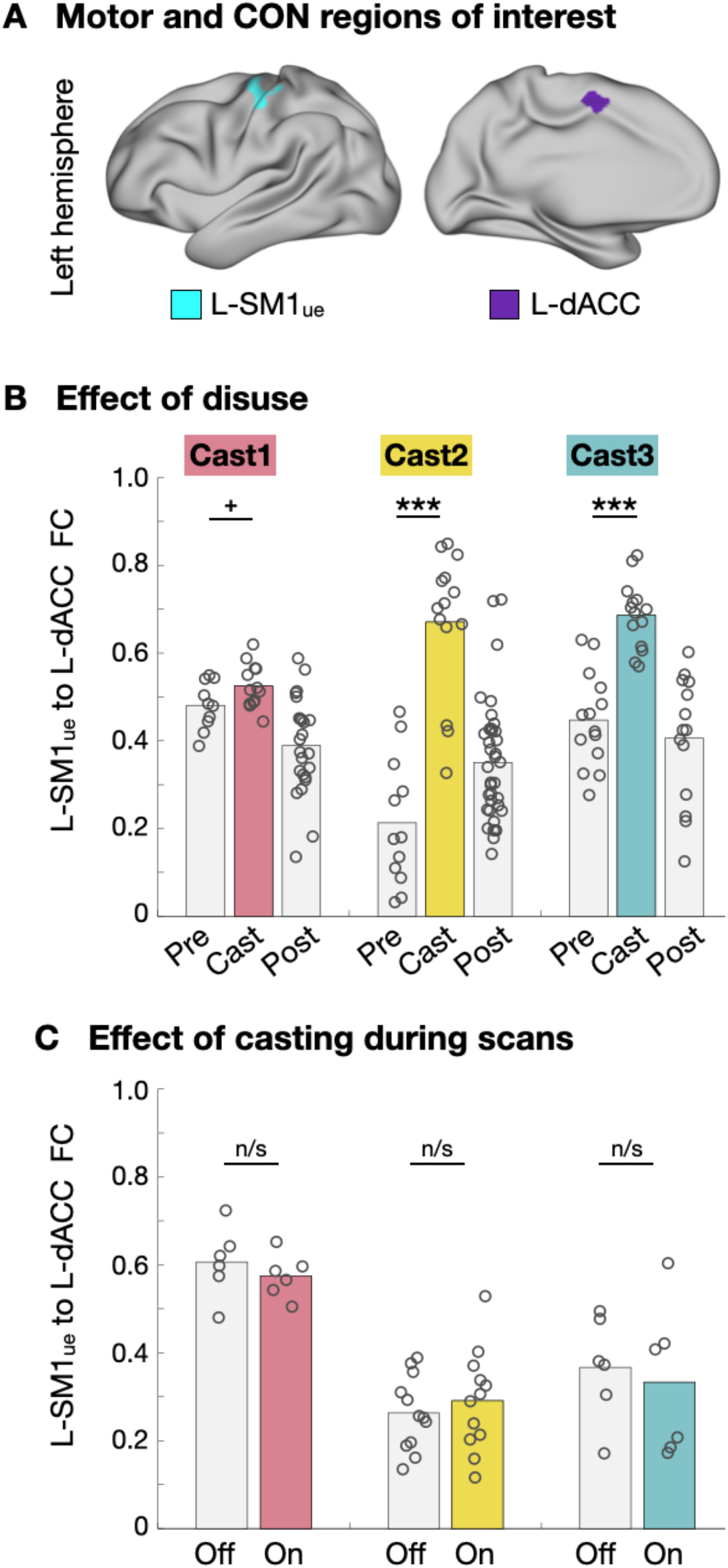
Connectivity between Disused Somatomotor Cortex and Cingulo-opercular Network during Disuse and Temporary Casting. (A) Example regions of interest (ROIs) in the left primary somatomotor cortex (L-SM1_ue_) and the left dorsal anterior cingulate cortex (L-dACC). (B) Functional connectivity (FC) measured between L-SM1_ue_ and L-dACC before (Pre), during (Cast) and after casting (Post). Two participants (Cast2 and Cast3) show a significant increase in FC during the cast period ^+^P < 0.1, *P < 0.05, ***P < 0.001. (C) FC measured between L-SM1_ue_ and L-dACC during both conditions of the control experiment: without casting (Off) and with temporary casts during scans (On). No changes in FC were observed.

**Fig. S5.**
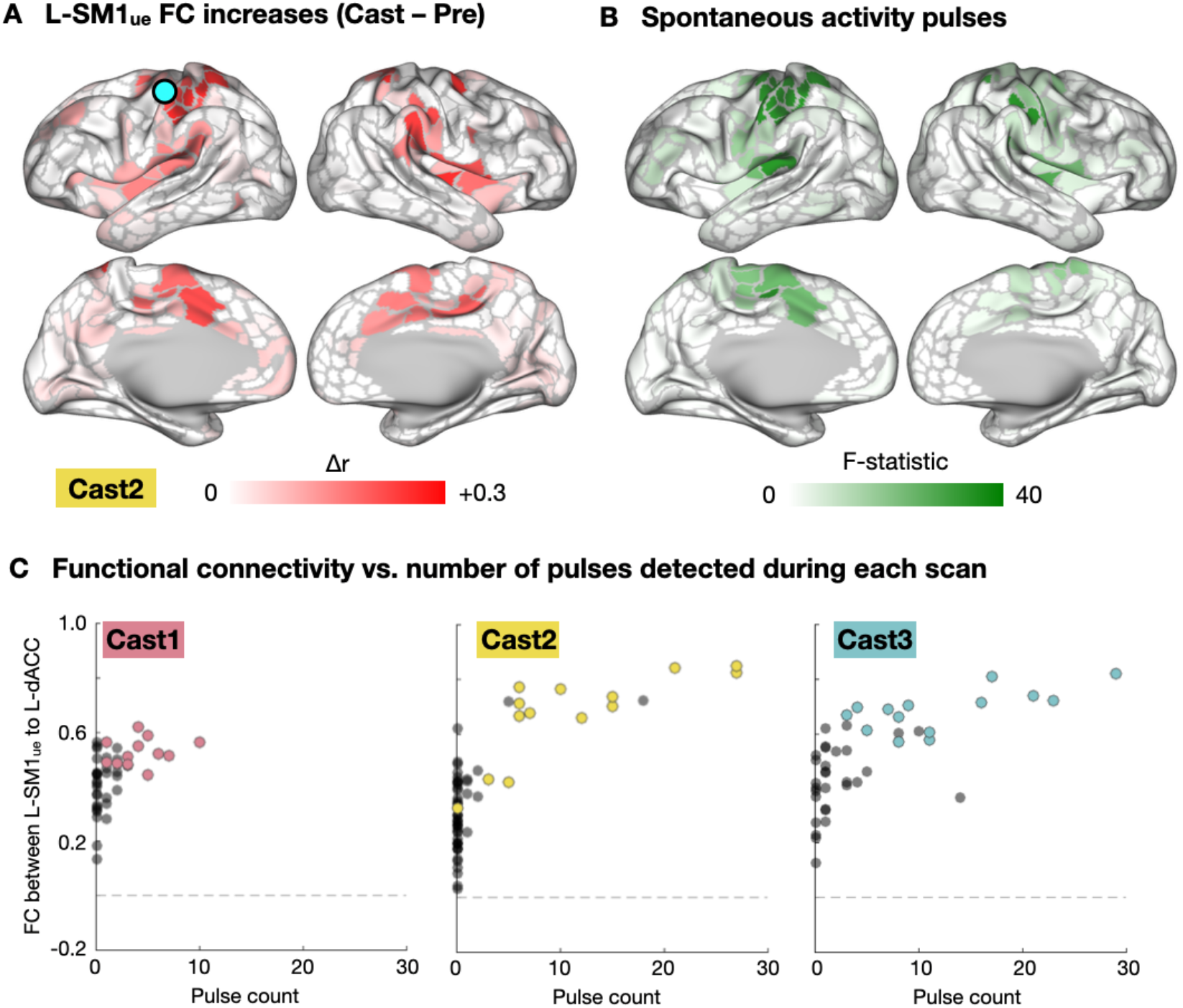
Relationship of Spontaneous Activity Pulses to Functional Connectivity (FC) Changes. (A) Increases in FC with the disused somatomotor cortex (L-SM1_ue_) during casting, repeated from Figure 2B for reference. (B) Analysis of variance (ANOVA) showing regions with spontaneous activity pulses. ANOVA F-statistic was correlated with FC change in each parcel (Cast1: r = 0.16, P < 0.001; Cast2: r = 0.29, P < 0.001; Cast3: r = 0.06, P = 0.12). (C) Relationship of FC between L-SM1_ue_ and the dorsal anterior cingulate cortex (dACC; cingulo-opercular network (CON)) vs. the number of pulses detected. Each dot represents one 30-minute scan. Colored dots represent scans during the cast period. For all participants, FC between L-SM1_ue_ and dACC was significantly correlated with the number of pulses detected (Cast1: r=0.74, P<0.001; Cast2: r=0.57, P<0.001; Cast3: r=0.73, P<0.001).

**Fig. S6.**
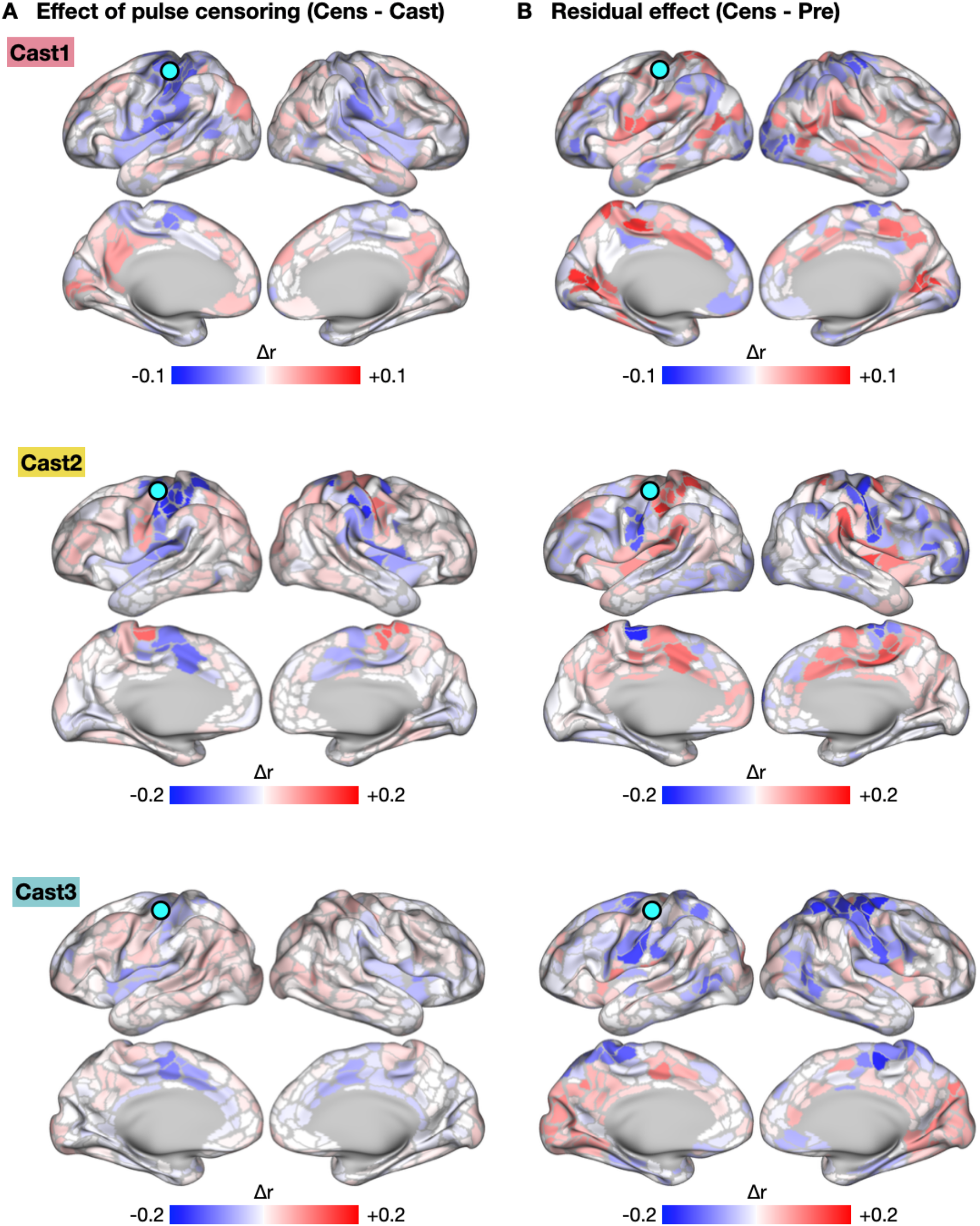
Functional Connectivity (FC) Changes Explained by Spontaneous Activity Pulses. (A) Subtraction map showing changes in L-SM1_ue_ FC due to pulse censoring. The spatial pattern of FC changes due to pulse censoring was negatively correlated with FC changes during casting (Cast1: r = −0.35, P < 0.001; Cast2: r = −0.90, P < 0.001; Cast3: r = −33, P < 0.001). (B) Subtraction map showing residual FC changes during casting after pulse censoring. The spatial pattern of residual FC changes was negatively correlated with the effect of pulse censoring (Cast1: spatial correlation, r = −0.23, P < 0.001; Cast2: r = −0.76, P < 0.001; Cast3: r = −0.18; P < 0.001).

**Fig. S7.**
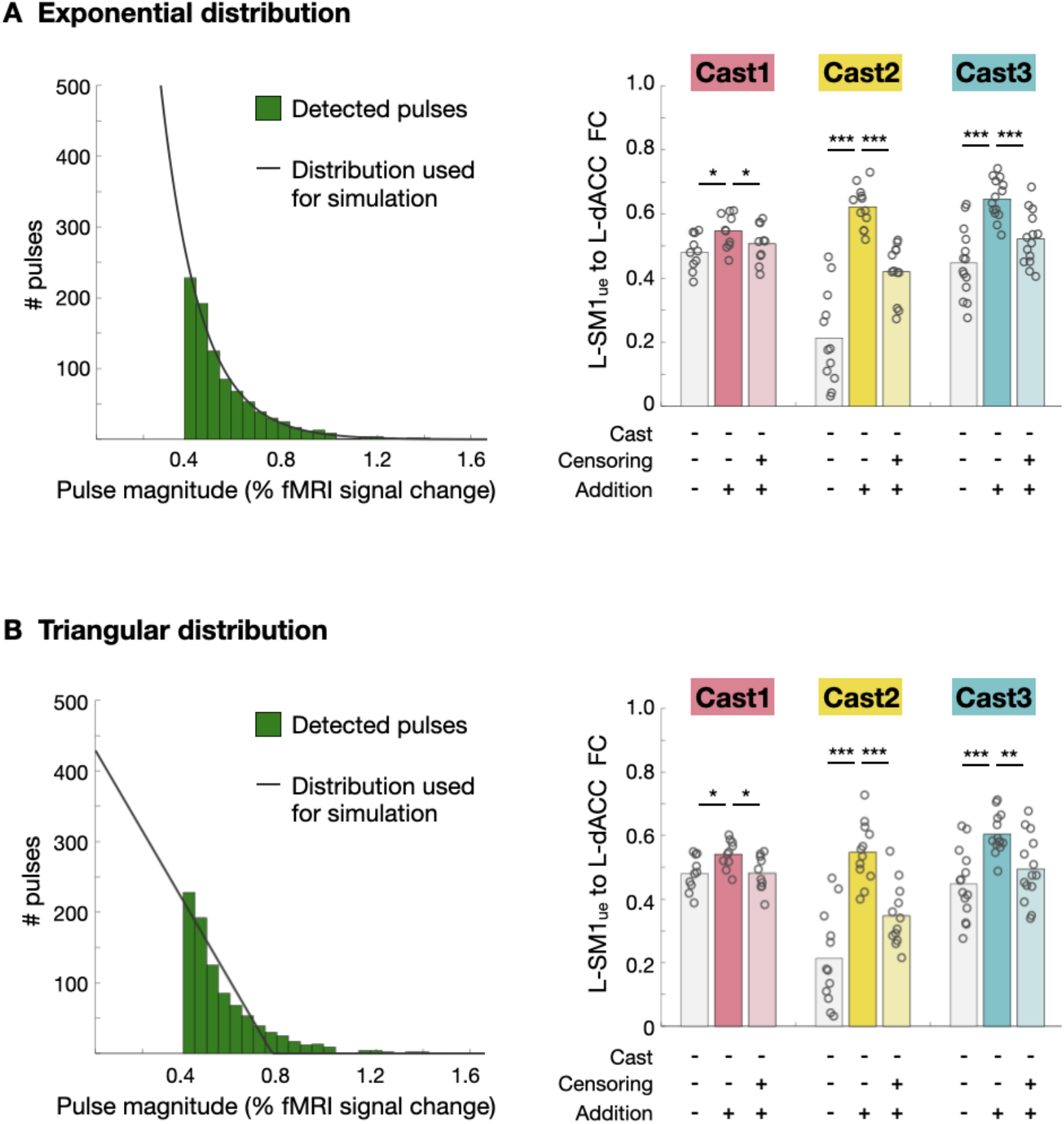
Alternative Distributions for Pulse Magnitudes. (A) Exponential distribution. *Left:* Histogram of pulse magnitudes (peak fMRI signal), pooled across all participants. An exponential distribution (black line) was fit to the data using a least-squares approach. *Right:* Adding simulated pulses, with magnitudes drawn from the exponential distribution, increased FC between L-SM1_ue_ and L-dACC. Censoring simulated pulses partially reduced FC increases. *P<0.05, ***P<0.001. (B) Triangular distribution. *Left:* Histogram of pulse magnitudes (peak fMRI signal), pooled across all participants. A linear distribution (black line) was fit to the data using a leastsquares approach. *Right:* Adding simulated pulses, with magnitudes drawn from the linear distribution, increased FC between L-SM1_ue_ and L-dACC. Censoring simulated pulses partially reduced FC increases. *P<0.05, **P<0.01, ***P<0.001.

**Fig. S8.**
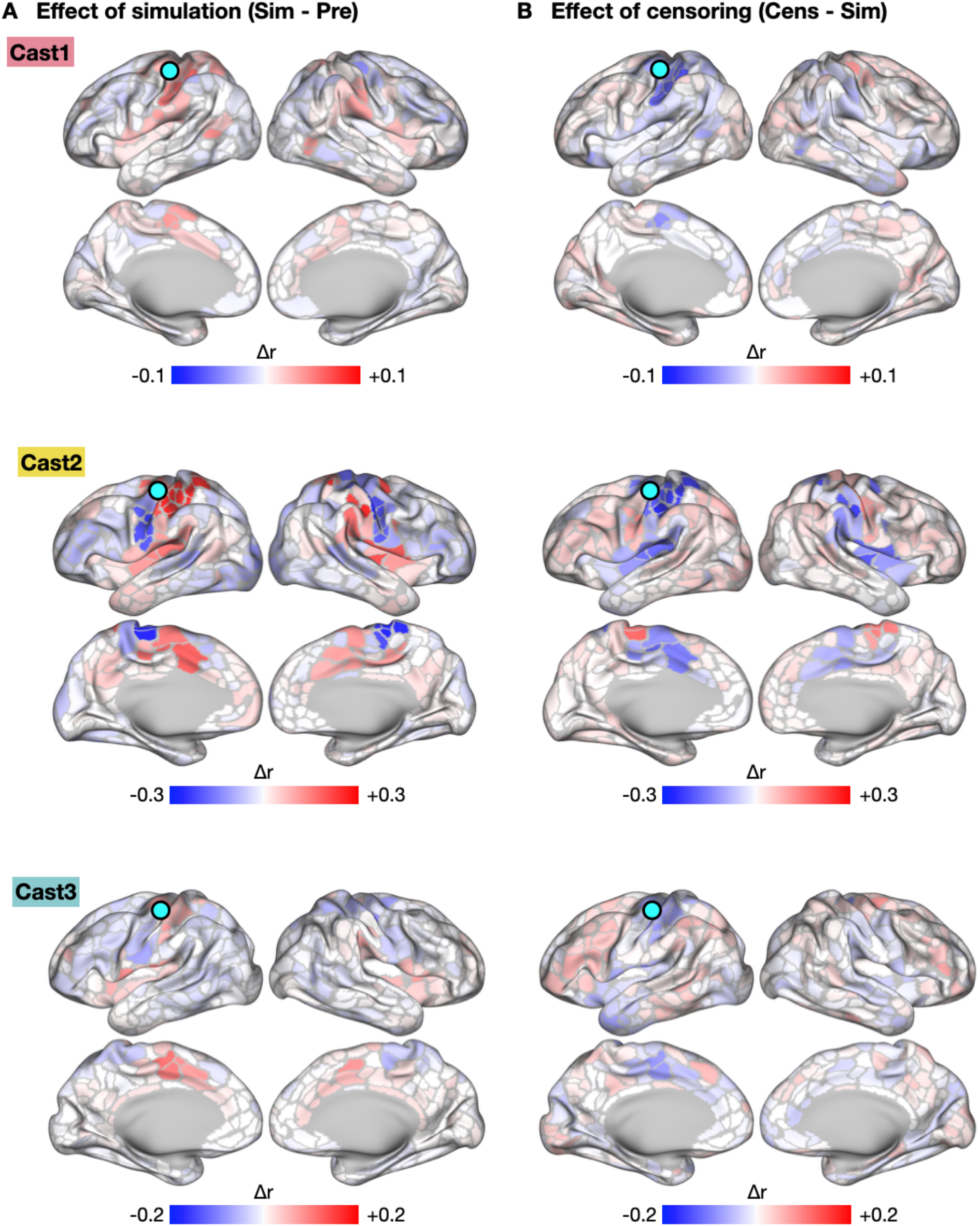
Functional Connectivity (FC) Changes Caused by Simulated Spontaneous Activity Pulses (All Participants). (A) Subtraction map showing changes in L-SM1_ue_ FC due to addition of simulated pulses. FC changes due to pulse addition closely matched FC changes during casting (Cast1: spatial correlation, r = 0.44, P < 0.001; Cast2: r = 0.95, P < 0.001; Cast3: r = 0.64, P < 0.001). (B) Subtraction map showing effect of censoring simulated pulses. Effects of censoring closely complemented the effect of pulse addition (Cast1: r = −0.71, P < 0.001; Cast2: r = −0.96, P < 0.001; Cast3: r = −0.69; P < 0.001).

**Fig. S9.**
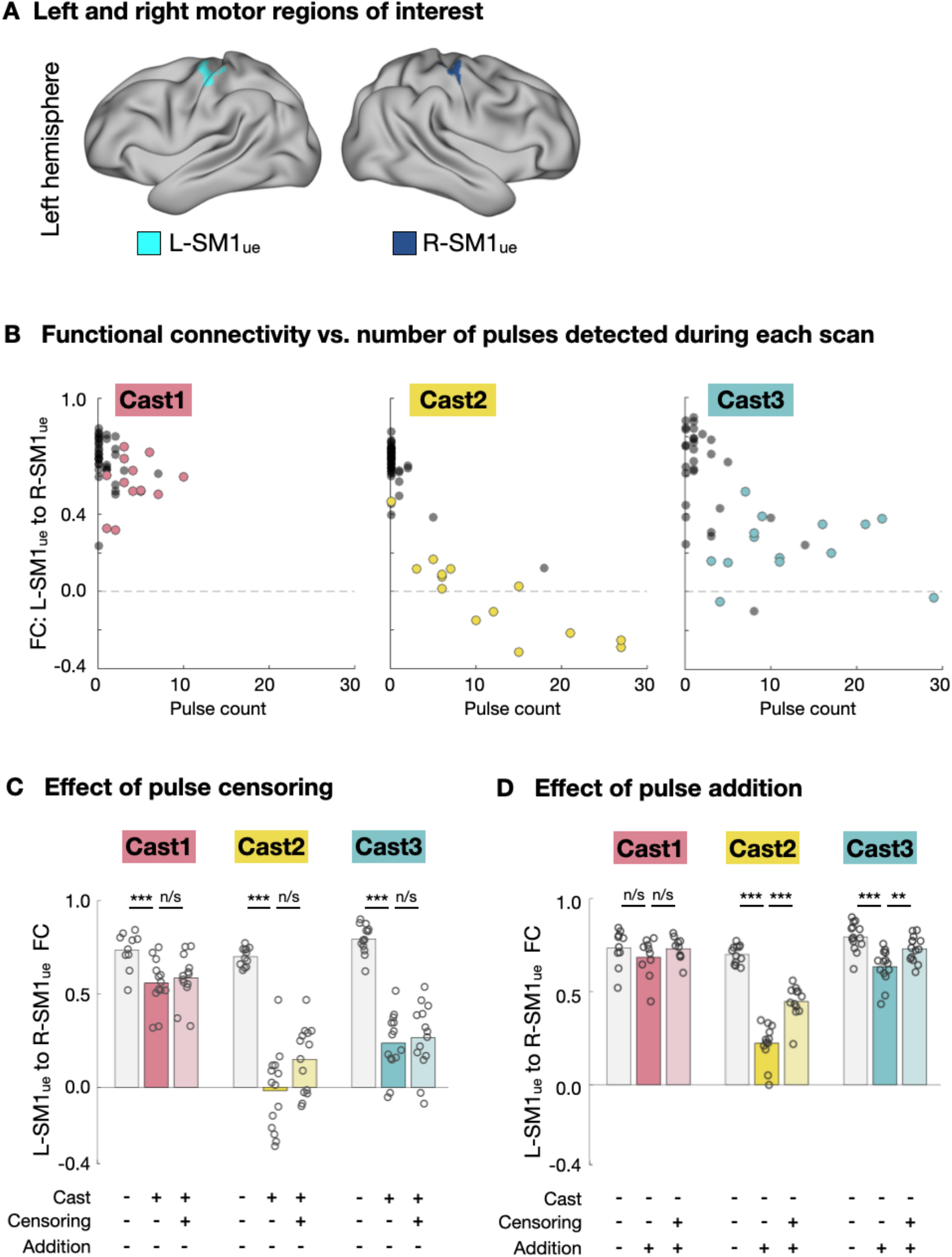
Functional Connectivity (FC) Decreases Were Not Explained by Spontaneous Activity Pulses. (A) Example regions of interest (ROIs) in the left and right primary somatomotor cortex (L-SM1_ue_ and R-SM1_ue_). (B) Relationship of FC between L-SM1_ue_ and R-SM1_ue_ vs. the number of pulses detected. Each dot represents one 30-minute scan. Colored dots represent scans during the cast period. For all participants, FC between L-SM1_ue_ and R-SM1_ue_ was significantly correlated with the number of pulses detected (Cast1: r=0.74, P<0.001; Cast2: r=0.57, P<0.001; Cast3: r=0.73, P<0.001). (C) Censoring pulses did not reduce the difference between Pre and Cast scans. ***P < 0.001. (D) Adding simulated pulses decreased FC between L-SM1_ue_ and R-SM1_ue_ in two participants (Cast2 and Cast3), but these decreases were much smaller than those observed during casting. ***P < 0.001. **P < 0.01.

